# The microstructure of laminin-111 compensates for dystroglycan loss in functional differentiation of mammary epithelial cells

**DOI:** 10.1101/531053

**Authors:** A Kent, N Mayer, JL Inman, C Hochman-Mendez, MJ Bissell, C Robertson

## Abstract

Laminin-111, an extracellular matrix (ECM) glycoprotein found in the basement membrane of mammary gland epithelia, is essential for lactation. In mammary epithelial cells, dystroglycan (Dg) is believed to be necessary for polymerization of laminin-111 into networks, thus we asked whether correct polymerization could compensate for Dg loss. Artificially polymerized laminin-111 and the laminin-glycoprotein mix Matrigel, both formed branching, spread networks with fractal dimensions from 1.7-1.8, whereas laminin-111 in media formed small aggregates without fractal properties (a fractal dimension of 2). In Dg knockout cells, either polymerized laminin-111 or Matrigel readily attached to the cell surface, whereas aggregated laminin-111 did not. In contrast, polymerized and aggregated laminin-111 bound similarly to Dg knock-ins. Both polymerized laminin-111 and Matrigel promoted cell rounding, clustering, formation of tight junctions, and expression of milk proteins, whereas aggregated Ln-1 did not attach to cells or promote functional differentiation.

**Highlights:**

- Laminin assembles into a fractal network when in presence of either the cell surface receptor dystroglycan or acidic glycoproteins or an acidic buffer.
- When this microstructure is recreated with an acidic treatment, laminin binds readily to dystroglycan null cells and induces functional differentiation of mammary epithelial cells.

## Introduction

Laminins in the basement membrane of the mammary gland play a powerful role in regulating the function of epithelial cells[1]. In particular, laminin-111 (Ln-1) is necessary for induction of milk protein expression in mammary epithelial cells (MEC) [2] and induction of proper architecture and quiescence [3, 4]. Furthermore, reversion of breast cancer cells to a quiescent phenotype requires appropriate cell-Ln-1 signaling [5], showing that induction of normal function is intimately coupled with suppression of malignancy [6]. Thus understanding the mechanisms by which Ln-1 signals to epithelial cells affords new insight into the processes in the breast that regulate normal behavior and suppress malignancy. Mammary epithelial cells have a range of receptors for Ln-1 including several integrins, dystroglycan and others and these different receptors appear to play different roles in mediating laminin binding and downstream signaling [1, 7]. Among the receptors for Ln-1, dystroglycan (Dg) plays a key role in mediating Ln-1 signaling [8, 9] and Dg is frequently lost or alternatively glycosylated in breast and other cancers [10, 11].

Dg is known to play two roles in laminin signaling: Dg links extracellular laminin to cytoskeletal actin [12] and Dg is essential for assembly of laminins in formation of basement membranes [13]. In mammary epithelia, evidence suggests that only the latter function is necessary. Loss of any of the intracellular domains of Dg does not perturb laminin binding or downstream signaling [8, 14], and conversely, loss of the extracellular alpha domain alone perturbs the cells ability to bind and respond to laminin [8, 15]. Furthermore, the extracellular domains of Dg grafted onto the transmembrane domain of a different protein is sufficient for Ln-1 binding to the cell surface and induction of milk proteins [8]. This suggests that Dg in the mammary gland does not directly transduce Ln-1 and instead acts through other mechanisms. However, the complex laminin-rich mixture of proteins secreted by EHS tumors (sold as Matrigel) does not show this same dependence on Dg for laminin anchorage [8], and deletion of Dg in the mammary gland does not appear to impair basement membrane formation in vivo [14]. Thus the mechanism by which Dg acts in laminin signaling remains unclear.

We directly tested the hypothesis that dystroglycan is necessary for MEC function because it polymerizes laminin into a basement membrane-like layer in the mammary gland. We used a recently developed method to artificially generate a hexagonal lattice of laminin proteins resembling the innermost layer of the basement membrane [16]. We then compared these networks structurally to the laminin networks in Matrigel, tested whether Dg knockout and knock in cells were sensitive to microstructural changes in purified laminin and evaluated whether synthesis of milk proteins and cell morphology changes could be induced by correctly structured laminin.

## Materials and Methods

### Laminin polymerization

Purified laminin-111 from EHS/Matrigel was purchased from Invitrogen (23017015). Growth factor reduced Matrigel was purchased from Corning (356230). For polyLM polymerization or control pH7 treatment, 100 µg/mL Ln-1 was mixed with 1 mM CaCl_2_ and 20 mM sodium acetate for pH4 or 1 mM CaCl_2_ and 20 mM Tris for pH7 for 30 minutes at 37°C as described in [16]. Treated polyLM laminin was then spun down at 2,000 rcf and treated pH7 laminin was spun down at 20,000 rcf to pellet laminins. The pellets were then resuspended in cell culture media for adsorption to glass slides or for addition to cells in culture. For laminin-nidogen-1 copolymers, we mixed laminin at 50 µg/mL with 50 µg/mL recombinant nidogen-1 (R&D systems 2570-Nd-050). Samples were immunostained as described below.

### Cell culture

Murine mammary DgKO and DgKI cells, originally derived from Dg conditional knockout mice [10] and Eph4 were cultured as previously described [8]. Briefly, cells were seeded at 8K cells/cm^2^ for DgKO or Eph4 or 5K cells/cm^2^ for DgKI to account for differences in growth rate and cultured in DMEM F12 with 2% FBS, 0.01 mg/mL Insulin (Sigma-Aldrich 1-1882), 0.005 µg/mL EGF (BD Bioscience 354001), 0.05 mg/mL Gentimicin (Thermo Fisher 15750060), and 0.05 mg/mL Normocin (InvivoGen ant-nr-1) in 37°C, humidified incubators at 5% CO_2_ tension. Cells were passed every 4-5 days by treatment with 0.25% Trypsin, neutralized with cell culture media (FBS), spun down at 115 rcf for 5 minutes and resuspended in media.

For laminin overlay procedures, cells were seeded at 10.5K cells/cm^2^ and grown for 2 days, then treated with Mg-Ln, polyLM or pH7 laminin in induction media comprised of DMEM/F12, 3 µg/mL prolactin (National Hormone and Peptide Program, Ovine PRL 10mg), 2.8 µM Hydrocortisone (Sigma-Aldrich H-0888 1G), and 5 *µ*g/mL Insulin (Sigma-Aldrich 1-1882). Cells were then cultured for 2 days before collection for analysis by immunostaining or western blot as described below.

### Immunostaining

Laminin polymers and cell cultures were fixed with 4% paraformaldehyde in, blocked with 10% goat serum plus Fab fragment (Jackson ImmunoResearch 715-007-003) to block Ln-Mg’s endogenous mouse antibodies, then stained with an anti-laminin antibody (Sigma L9393) or anti-ZO-1 (Invitrogen 61-7300) in block solution overnight at 4°C, followed by Alexafluor conjugated secondary antibodies (Invitrogen A-11034) and DAPI (4’,6-Diamidino-2-Phenylindole, Dihydrochloride) when appropriate.

### Microscopy and Image Analysis

Widefield microscopy was performed on an upright Zeiss Axioimager and confocal microscopy was performed on a Zeiss LSM 710 scope with a 63X/1.4NA objective. Pinhole was set to one Airy unit. Images of laminin networks were processed to measure the box counting dimension in MATLAB (Mathworks, Natick). Cross correlation and autocorrelation analyses was performed in MATLAB as described previously [17-19].

### Scanning Electron Microscopy

Laminin matrices generated at pH4 or pH7 were spun down and resuspended in cell culture media (pH 7.4) and allowed to adsorb onto silica wafers for 90 minutes. Matrices were fixed in 2.5% glutaraldehyde in 0.1M sodium cacodylate buffer, pH 7.4, washed three times with 0.1M sodium cacodylate buffer, pH 7.2, post-fixed 1 hour in 1% osmium tetroxide in 0.1M sodium cacodylate buffer, pH 7.2, washed three times in 0.1M sodium cacodylate buffer, pH 7.2, dehydrated through increasing concentrations of ethanol and sputter coated with a thin layer of gold. Samples were imaged on the Hitachi S-5000 scanning electron microscope.

### Western Blotting

For western blotting, cells were harvested in RIPA buffer with protease inhibitor (Calbiochem 539 131-10VL) and phosphatase inhibitors 1 and 2 (Sigma-Aldrich P2850-5ML and P5726-5ML, respectively) on ice and then passed through a 29G insulin needle and then spun down for 20 mins at 21,000 rcf. Supernatant was kept and the protein concentration was determined using a Lowry Assay (DC Protein Assay Bio Rad 500-0115). Proteins were loaded on 4-20%Tris-Gly gradient gels (invitrogen XP04200BOX) then transferred to nitrocellulose membranes with a dry transfer system (Invitrogen iBlot Blotting System and Gel Transfer Stacks Invitrogen IB301001). Membranes were stained with anti-laminin (Sigma L9393), rabbit polyclonal anti-β casein (developed by Caroline Kaetzel [20]), anti-lamin B1 (Abeam abl6048) or anti-α-tubulin (Santa Cruz 32293).

### Statistical analyses

ANOVA analyses and Tukey’s posthoc tests were performed in MATLAB with a significance level of p<0.05.

## Results

### PolyLM laminin networks are structurally similar to laminin-glycoprotein mixes

To determine why Dg is necessary for Ln-1 signaling in epithelial cells, we cultured Dg knockout (DgKO) mouse MEC and this line with Dg expression was restored by knock-in (DgKI) (created in [8]) - in the presence of the lactogenic hormones and an overlay of purified Ln-1 (as described in [8]). We observed that Ln-1 on the surface of DgKI cells formed web-like networks, whereas in acellular regions Ln-1 formed small aggregates (Fig. 1A and insert). In DgKO cultures, similar aggregates were observed in acellular regions, but minimal Ln-1 was observed on the surface of cells (Fig. 1B and insert).

**Fig. 1:**
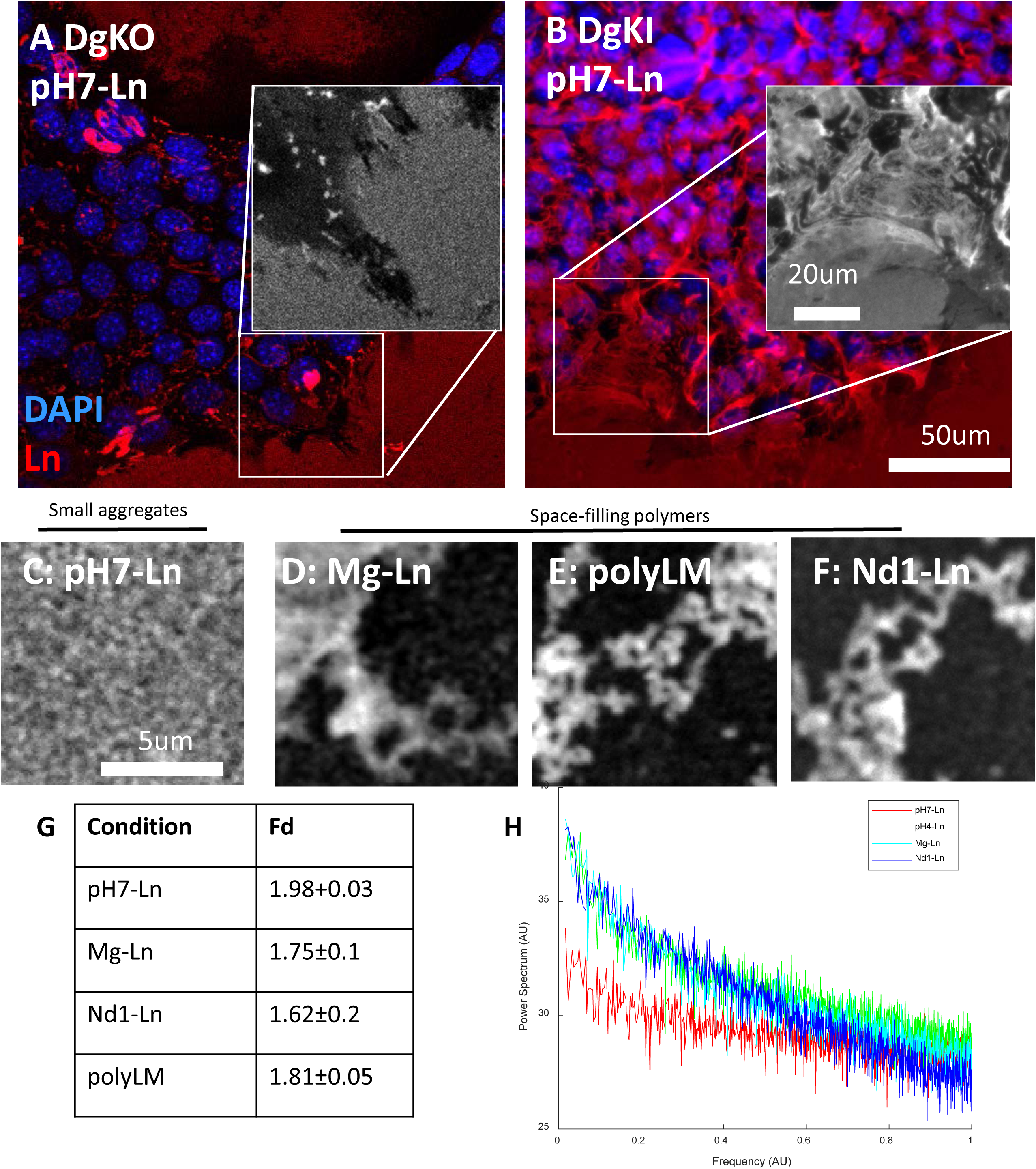
Fractal laminin polymerization occurs naturally on DgKI cells and artificially in acidic conditions. A. Immunofluorescence of purified Ln-1 on DgKO at pH7 shows that laminin fails to adhere to cells in the absence of Dg. *Inset:* almost no laminin is observed on the cell surface and laminin adhered to glass in acellular regions forms globules B. Immunofluorescence of purified Ln-1 on DgKI cells at pH7 shows that laminin adheres and forms a lacy network. *Inset:* laminin network on the cell surface shows a different morphology compared to on the glass, C. Laminin in a neutral pH7 buffer forms small globular aggregates D. Laminin network in the laminin-glycoprotein mix Matrigel ™ (Mg-Ln) forms web-like networks. E. Laminin in an acidic pH4 buffer (polyLM) forms a web-like network similar to that of Mg-Ln F. Laminin in the presence of the acidic glycoprotein Nidogen (Nd1-Ln) forms a polymerized laminin network similar to that of Mg-Ln G. Fractal dimensions calculated by box counting dimension for the four tested conditions. Mg-Ln, polyLM and Nd1-Ln all have fractal dimensions around 1.7 which is typical of diffusion limited aggregates, whereas pH7-Ln has a distinct, non-fractal dimension of 2. H. Power spectrum for images of all four acellular laminin conditions tested. Note the similarities for Mg-Ln, Nd1-Ln and polyLM despite the differences in synthesis methods.

We compared the microstructure of purified laminin aggregated in media, to the laminin network formed in the laminin-glycoprotein mix Matrigel, to polymerized purified laminin [21-23]. Laminin adsorbed to glass in neutral buffered media (pH7-Ln) formed small aggregates (Fig. 1C), whereas polyLM, and Mg-Ln conditions formed branched, space filling polymers (Fig. 1D&E). To evaluate why Mg-Ln in neutral buffered media showed such a different network structure than purified Ln-1 in the same conditions, we mixed Ln-1 with recombinant nidogen-1 and observed network formation similar to Mg-Ln and polyLM (Fig. 1F). Interestingly, switching from phosphate buffered saline to tris buffered saline destroyed the network structure of Ln for each of these conditions, suggesting that pH may underlie network assembly (Supplemental Fig. 1).

As previous work has demonstrated that polymerized laminin has fractal properties [21], we quantitatively compared network morphology, using the both box counting algorithm and power spectrum analysis to estimate the fractal dimension of these images. We found that polyLM, Nd1-Ln, and Mg-Ln had fractal dimensions of approximately 1.7 consistent with previous work [21] (Fig. 1E), whereas pH7-Ln had a dimension of approximately 2. Power spectrum analysis, an alternative method to measure the microstructure of fractal images without thresholding [18], likewise showed similar properties for polyLM, Nd1-Ln and Mg and dramatically different measures for pH7-Ln (Fig. 1H). Thus, the Ln-1 networks formed in Mg-Ln, Nd1-Ln and polyLM were structurally similar, whereas pH7-Ln had markedly different morphological properties.

To confirm these findings using higher resolution imaging, we collected pH7-Ln, polyLM, and Mg-Ln onto silica wafers, then processed these samples for scanning electron microscopy (Fig. 2A-C). We observed a lacy network structure in polyLM and Mg-Ln which was absent in pH7-Ln conditions suggesting that Mg-Ln and polyLM show similar organized network structure (Fig. 2D).

**Fig. 2:**
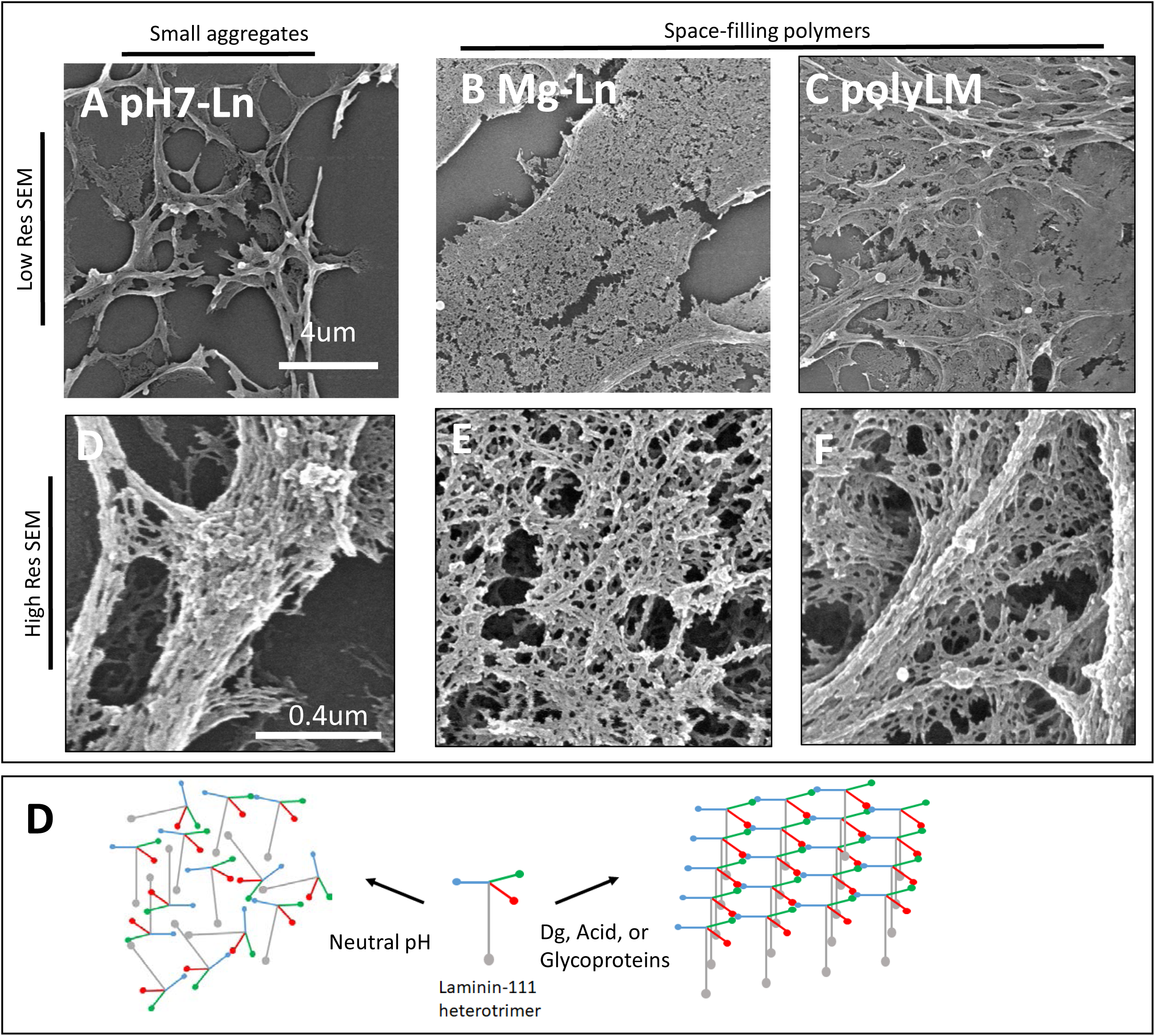
Electron microscopy characterization of laminin networks shows similarity between Mg-LN and polyLM at fine scales. A-C. Scanning electron microscopy of pH7-Ln, Mg-Ln and polyLM at 2 resolution scales. Note that at high resolution both Mg-Ln and polyLM share a lacy network structure, whereas pH7-Ln forms globules D. Cartoon of theorized network structures. In neutral pH laminin assembles randomly, resulting in globules, whereas in the presence of Dg, low pH, or glycoproteins, laminin assembles into networks with a hexagonal lattice at the atomic scale and fractal networks at the ultrastructural scale.

### The microstructure of laminin alters its binding to DgKO but not to DgKI cells

Next, we compared whether DgKO and DgKI mouse MEC bound polyLM and pH7-Ln using Mg-Ln as a positive control. PolyLM, pH7-Ln and Mg-Ln all adhered to the surface of DgKI cells (Fig. 3A) and similar amounts of polyLM and pH7-Ln bound to DgKI cells by both western blot or immunofluorescence (p>0.5) (Fig. 3A). In DgKO cells, polyLM and Mg-Ln both adhered to cells readily (Fig. 2B), and the amount of Ln-1 bound to cells in polyLM was significantly higher than in pH7-Ln conditions.

**Fig. 3:**
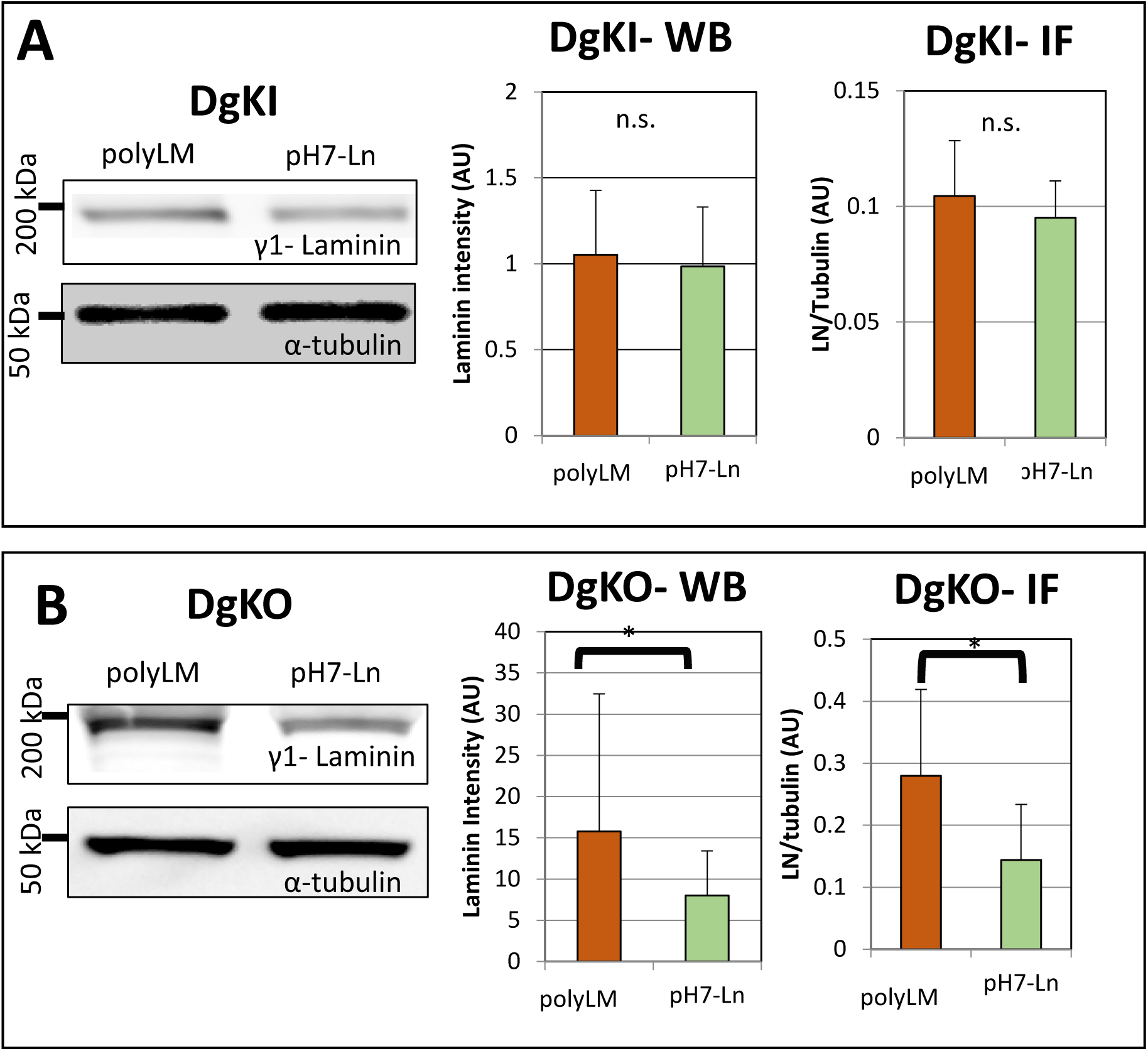
Laminin polymerization affects its adherence to DgKO cells. A. Both polyLM and pH7-Ln adhere to DgKI cells. Western blots for Ln-1 (tubulin loading control) show that artificially polymerizing laminin in acidic buffer or leaving it non-polymerized in a neutral buffer does not affect its ability to bind to DgKI cells. Quantification of western blot and immunofluorescence show no significant difference between polyLM and pH7-Ln adherence to DgKI cells. B. PolyLM but not pH7-Ln adheres to DgKO cells. Western Blot for laminin (tubulin loading control) show that artificially polymerizing laminin in an acidic buffer (polyLM) causes more Ln1 to bind to DgKO cells relative to the non-polymerized laminin in a neutral buffer. Quantification of western blots and immunofluorescence confirm a significant increase of polyLM adherence to DgKO cells relative to pH7-Ln.

To demonstrate that the difference in Ln-1 levels observed was due to anchorage of exogenously provided laminin and not due to differences in Ln-1 production, we repeated these experiments with fluorescently labeled Ln-1 (Fig. 4) and observed a low level of endogenous Ln-1 in DgKO cultures (Fig. 4A&C) in pH7-Ln conditions, and a mix of exogenous Ln-1 and some endogenous production in polyLM conditions (Fig. 4B&D). This shows that engagement of Ln-1 with the surface of cells requires either Dg or glycoproteins or correctly polymerized Ln-1.

**Fig. 4:**
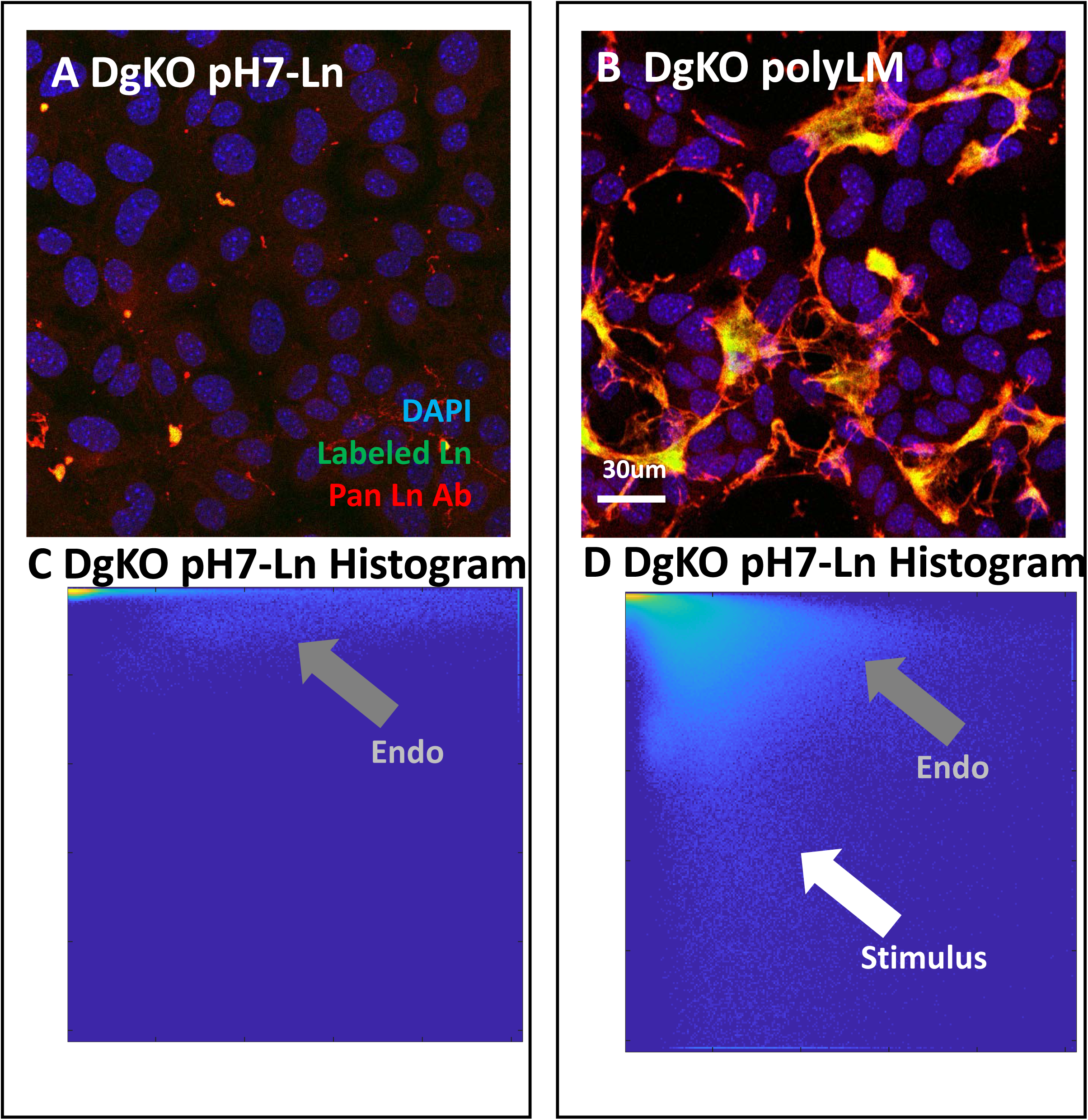
Laminin polymerization induces expression of endogenous laminin. A-B. DgKO cells were treated with fluorescently labeled laminin (A-fluorescent pH7-Ln, B-fluorescent polyLM) then fixed and stained using an anti-laminin antibody. We generated 2d histograms of intensity of the fluorescent tag and antibody staining for these images C. In pH7-Ln conditions antibody staining shows low levels of endogenously produced laminins but essentially no labeled exogenous Ln-1 (gray arrow). D. In polyLM conditions, we observe both exogenous Ln-1 (white arrow-high antibody stain, and high fluorescent label) and formation of endogenous laminins (gray arrow-high antibody staining, no fluorescent tag).

### PolyLM induces functional differentiation of Dystroglycan Knockout MEC

Given that polymerizing Ln-1 restored its adhesion to cells, we next asked in DgKO MEC if Ln-1 microstructure could induce differentiation by four metrics-induction of β-casein expression, colony morphology, tight junction formation, and reduction of cytoplasmic actin and actin stress fibers.

We observed significantly higher expression of β-casein in polyLM conditions compared to pH7-Ln (Fig. 5A&B). As Mg-Ln was more efficient in binding to cells at similar protein doses, we compared laminin binding and morphological changes with increasing doses of either Mg-Ln, polyLM, or pH7-Ln (Fig. 5C and 5D-I). We observed increased laminin staining by quantitative image analysis in both the Mg-Ln and polyLM (Fig. 5J) conditions but not in pH7-Ln conditions. Notably, polyLM at 800 μg/mL and Mg-Ln at 50μg/mL were not statistically different with respect to Ln-1 anchorage or β-casein expression.

**Fig. 5:**
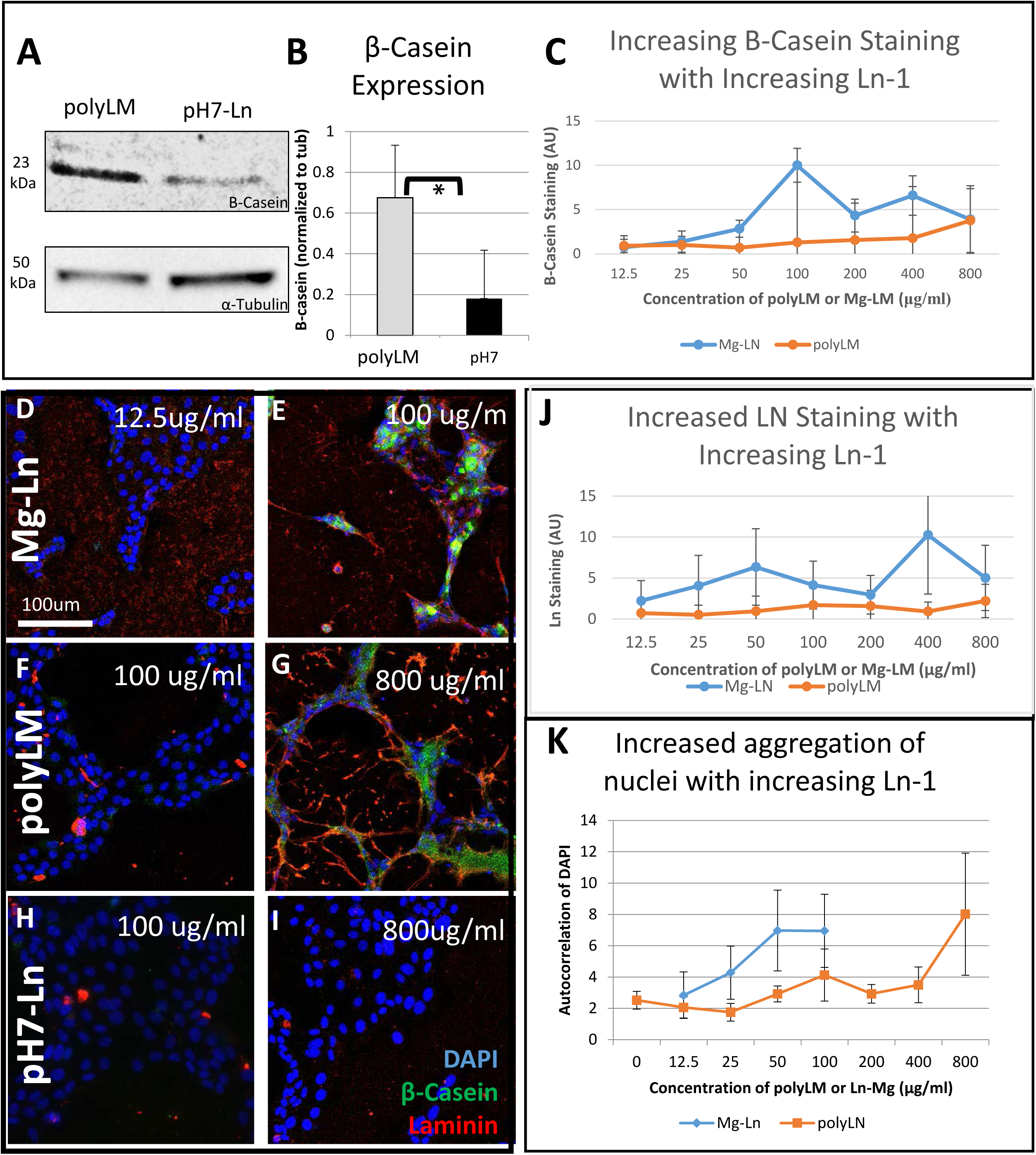
Laminin polymerization affects milk protein expression and cell morphology. A. βcasein expression is higher in polyLM than pH7-Ln stimulated DgKO cells. B. Quantification of western blots shows significantly higher β-casein expression in polyLM conditions C. With increasing doses of either polyLM or Mg-Ln increasing expression of β-casein is observed by immunofluorescent staining and quantitative image processing. Notably β-casein expression in Mg-Ln at 50ug/ml and polyLM at 800ug/ml is not statistically different. D-l. Cell colony morphology for Mg-Ln at low (D) and for polyLM at low (F) shows flat colonies whereas at high doses both Mg-Ln (E) and polyLM (G) form web like networks. pH7-Ln at both low and high dose (H-l) shows flat spread colonies with no laminin binding. J Quantification of Ln-1 staining with increasing stimulus of either Mg-Ln or polyLM K. Autocorrelation analysis indicates increased clustering of nuclei with increased doses of either Mg-Ln or polyLM. Mg-Ln at 50 or 100 ug/ml and polyLM at 800ug/ml are statistically different than pH7-Ln at 800ug/ml.

Concomitant with induction of β-casein expression, we observed a colony morphology change from relatively flat, spread colonies (Fig. 5D&F) to formation of tubular colonies with increasing doses of both polyLM and Mg-Ln (Fig. 5E&G). These morphological changes were not observed even the highest tested dose (800ug/ml) of pH7-Ln (Fig. 5I). To quantify these changes in cell colony morphology, we calculated the autocorrelation of the images of nuclei [17, 18] (Fig. 5K) then compared across Ln treatment and concentration. We observed that cells in Mg-Ln at 50 or 100ug/ml or polyLM at 800ug/ml showed statistically higher aggregation compared to pH7-Ln at 800ug/ml by 2-way ANOVA and Tukey’s post hoc test (Fig. 5K).

Given this formation of tubular cell colonies, we next investigated cell junction and cytoskeletal organization. In both the polyLM and Mg-Ln conditions, we observed continuous apical tight junctions containing both zonula occludens-1 (ZO-1) and filamentous actin (Fig. 6A&B), whereas interrupted ZO-1 staining was observed in either pH7-Ln or control no-Ln-1 conditions (Fig. 6C&D-indicated by gray arrows). Cross correlation analysis and 1-way ANOVA demonstrated that ZO-1 and actin were more likely to be colocalized in Mg-Ln or polyLM than in either pH7-Ln or no-Ln-1 controls (Fig. 6E), and association of ZO-1 and actin in pH7-Ln and control conditions was not statistically different.

**Fig. 6:**
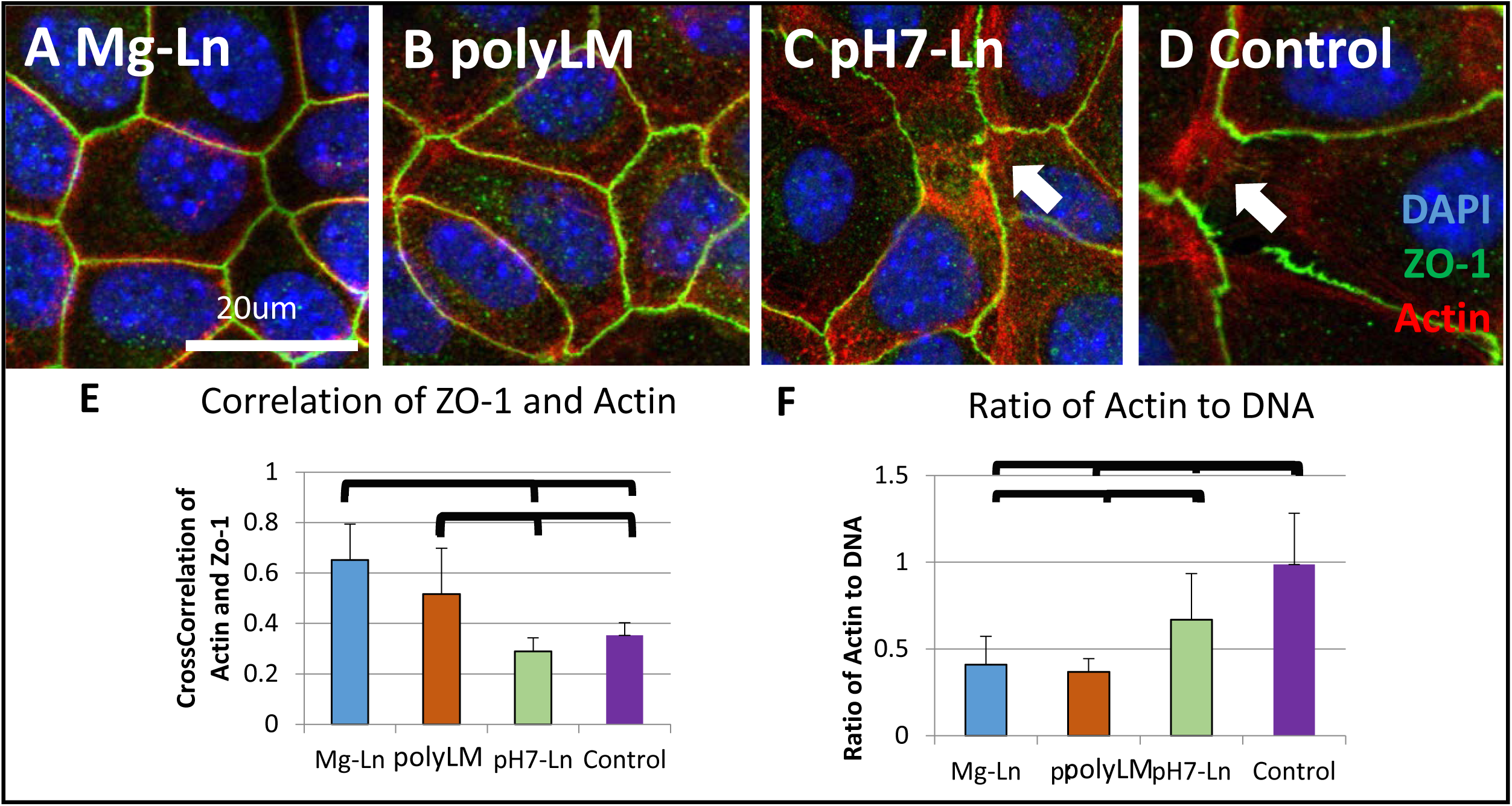
Laminin polymerization affects cell morphology. A-D. Cell-cell junctions vary with laminin treatment. In Mg-Ln conditions (A), apical tight junctions were well formed and occasional lumens were observed. In polyLM (B), colonies tended to be less well organized, but tight junctions between cells were well organized. In contrast, in pH7-Ln (C) and no-Ln-1 control conditions (D), gaps between cells and abundant actin stress fibers were observed. E. In Mg-Ln and polyLM conditions, actin and ZO-1 were more likely to be colocalized by cross correlation analysis than in pH7-Ln or control conditions.

In polyLM or Mg-Ln, the majority of actin in the cell body colocalized with ZO-1 in the cell-cell junctions and almost no stress fiber formation was observed. In contrast in both pH-7-Ln and control conditions MEC formed abundant stress fibers at the basal surface (Fig. 6C&D). We observed a higher ratio of filamentous actin to DNA in either pH7-Ln or control conditions relative to polyLM or Mg-LN (Fig. 6F). Furthermore, we determined that cortical actin and laminin were anticorrelated in both Ln-Mg and polyLM conditions (Supplemental Fig. 2).

## Discussion

In this work, we demonstrated that mammary epithelial cells depend on Ln-1 microstructure and that this change in anchorage regulates milk protein expression and cell shape. We found that multiple mechanisms give rise to similar Ln-1 microstructures. Ln-1 polymerized into a fractal network in the presence of Dg, acidic proteoglycans such as those found in Matrigel, or acidic buffered calcium solutions, suggesting these mechanisms can compensate for each other. We found that polymerized Ln-1 formed a coherent coat on the cell surface and induced cell shape changes and β-casein expression. The complex mixture of laminins and glycoproteins in Mg-Ln was more efficient in inducing these changes than polyLM.

That Dg is dispensable for induction of β-casein is surprising. In myocytes, Dg is necessary sarcomere anchorage and organization and thus prevents a form of muscular dystrophy [24, 25]. In contrast we found that cortical actin and Ln-1 were anti-correlated in MEC, with less cortical actin in regions where Ln-1 bound to the cell surface, demonstrating tissue specific differences in how Ln-1 is sensed (Fig. S2). Nonetheless, the frequent loss or aberrant glycosylation of Dg in breast cancers suggests that it plays an important role in healthy breast epithelia [8].

Similar microstructures were created by both cell surface Dg and basement membrane proteoglycans, suggesting compensation between these mechanisms for polymerizing Ln-1 in some cases. Indeed, both Dg and Nidogen-1 knockout mice show essentially normal basement membrane formation and maintenance in epithelial tissues [26-29], suggesting that the compensation we observed in vitro may occur in vivo. We have demonstrated that polymerized Ln-1 can compensate for Dg loss, raising the question of how Ln-1 microstructure alters avidity of Ln-1 for cells.

Ln-1 is a trimeric, cross-shaped protein with different adhesive domains on the ends of the three short arms and one long arm thus the polymerization state may relate to the relative display of these different domains [30]. A number of laminin receptors bind only to the globular domains on the long arm or only to the short arms [31, 32]. At neutral pH in the presence of calcium, Ln-1 forms small aggregates which we believe is due to mixed short arm-short arm and short arm-long arm binding [21]. In contrast lowering the pH appears to preferentially charge the long arm globular domains of laminin, resulting in electrostatic repulsion between long arms and increased probability of short arm-short arm binding [21]. As a result, polymerization in acidic conditions appears to increase long arm display.

Previous work suggests that nidogen-1 and other glycoproteins crosslink specific domains on the long arm of Ln-1 [22]. It is therefore surprising that Mg-Ln or Nd1-Ln polymerized similarly to polyLM. As reported previously [33], we found that Ln-1 polymerized into a self-similar network with fractal dimension of ~1.7, characteristic of diffusion limited aggregation. Indeed, previous work shows that polyLM polymerization follows a nucleation and growth schema [21] which define diffusion limited aggregation phenomena. This suggests that laminin-glycoprotein mixes undergo similar aggregation processes for assembly. Indeed, we found that stronger buffer systems reduced the ability of nidogen-1 to promote polymerization suggesting that the electrostatic effects of negatively charged glycoproteins on laminin may be important (data not shown).

In all culture conditions tested, Mg-Ln was more efficient than purified Ln-1 alone in binding to MEC and inducing functional differentiation, suggesting that the additional protein components in Mg-Ln play some role in cell signaling and/or Ln-1 organization. Here we found that at similar total protein concentrations Mg-Ln formed coherent Ln-1 coats whereas polyLM formed patches. This finding is supported by previous work showing that while Nidogen-1 by itself does not induce β-casein expression in mammary cells, Nd1-Ln mixes are more effective than Ln-1 alone [34, 35].

We postulate that this enhancement is due to reinforcement of Ln-1 networks by other proteins or glycoproteins. Previous work has demonstrated that while Ln-1 is sufficient for initial formation of basement membranes, reinforcement with other proteins such as collagen IV is required for basement membrane stability [36]. Formation of apical polarity in epithelial cells has been linked previously to collagen IV reinforcement [37]. In agreement, we observed occasional formation of lumens in Mg-Ln conditions, whereas we did not in polyLM. This demonstrates that while polymerized Ln-1 is sufficient for induction of β-casein, it may not be sufficient for induction of all mammary epithelial specific phenotypes.

## Conclusions

In summary, we conclude that microstructure of Ln-1 networks in the basement membrane regulates epithelial cell function. We found that polymerized Ln-1 could come from either cell surface dystroglycan or basement membrane glycoproteins. In DgKO cells, polymerized laminin could drive cell morphology changes, tight junction assembly and milk protein expression.

## Acknowledgements

This work funded by the DOD CDMRP Breast Cancer Research Program BC133875, the L’Oreal USA for Women in Science Program, and Lawrence Livermore National Lab LDRD 18-ERD-062 to C.R., NIH R01CA064786, DOD Breast Cancer Research Program and the Breast Cancer Research Foundation (to MJB). Thanks to John Muschler for his kind gift of the DgKO and DgKI cells. Thank you to the staff at the University of California Berkeley Electron Microscope Laboratory for advice and assistance in electron microscopy sample preparation and data collection. Thanks to Cerise Bennett, Gita Rohanitazangi, Simone Skiba, Kate Thi, and Josephine Wu for their help with the work and to Monica Moya for her help with editing.

This work was performed under the auspices of the U.S. Department of Energy by Lawrence Livermore National Laboratory under Contract DE-AC52-07NA27344. LLNL-JRNL-766544

## Supplemental Figures

**Supplemental Figure 1:**
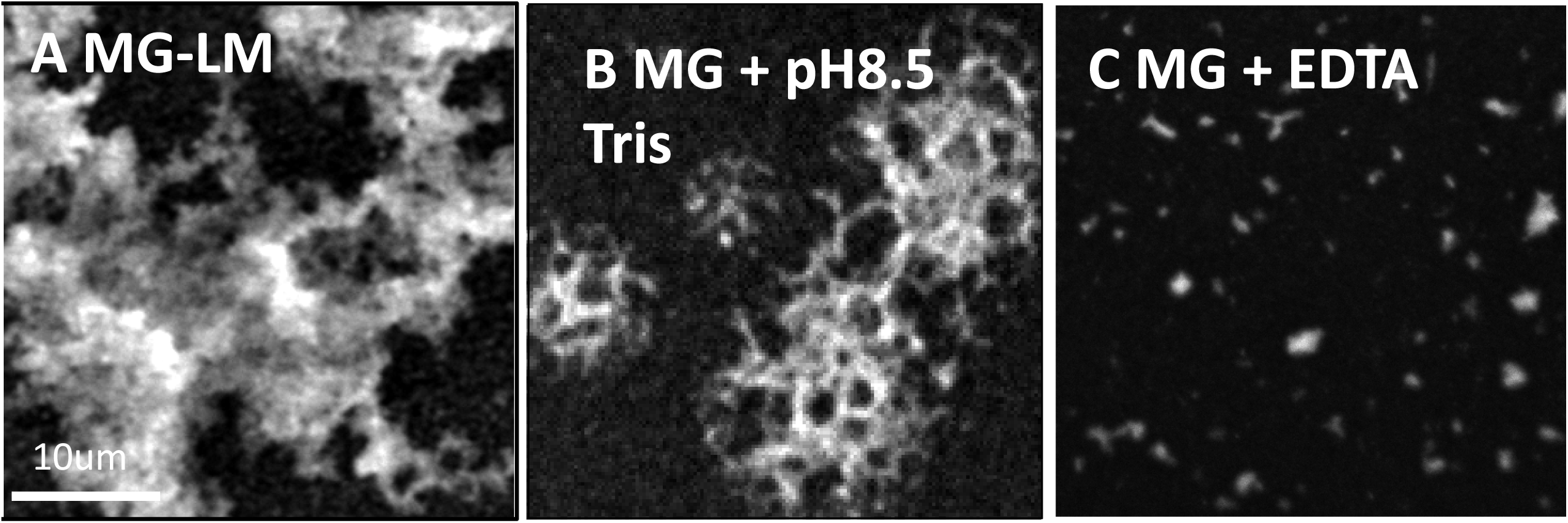
Organization of Mg-Ln depends on pH. A: Mg-Ln in neutral buffer forms a lacy network. B: when this is repeated at pH8.5, network structure is partially disrupted. C: Addition of the chelator EDTA completely destroys Mg-Ln polymerization.

**Supplemental Figure 2:**
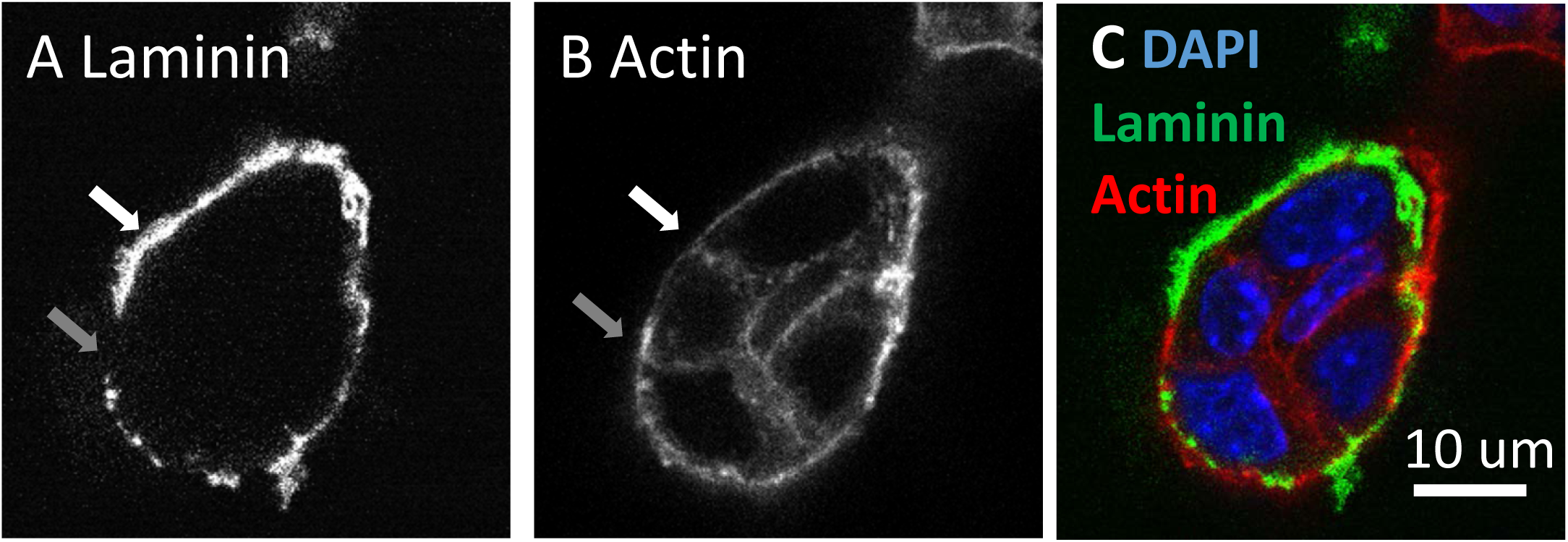
Laminin and cortical actin are anticorrelated. A: MEC stained for laminin shows concentration of laminin at the surface of cells. B: MEC stained with phalloidin for fibrillar actin shows concentration of actin at the lateral cell surface and in between cells. Gray area indicates an area of low laminin and abundant actin, white arrow shows a region of high laminin and low actin C: Merged image with DAPI in blue, laminin in green and actin in red. Note that at the cell surface, laminin and actin are anticorrelated.

